# Conserved Architecture of Brain Transcriptome Changes between Alzheimer’s Disease and Progressive Supranuclear Palsy in Pathologically Affected and Unaffected Regions

**DOI:** 10.1101/2021.01.18.426999

**Authors:** Xue Wang, Mariet Allen, Joseph S. Reddy, Minerva M. Carrasquillo, Yan W. Asmann, Cory Funk, Thuy T. Nguyen, Kimberly G. Malphrus, Steven G. Younkin, Dennis W. Dickson, Nathan D. Price, Todd E. Golde, Nilüfer Ertekin-Taner

## Abstract

We identify a striking correlation in the directionality and magnitude of gene expression changes in brain transcriptomes between Alzheimer’s disease (AD) and Progressive Supranuclear Palsy (PSP). Further, the transcriptome architecture in AD and PSP is highly conserved between the temporal and cerebellar cortices, indicating highly similar transcriptional changes occur in pathologically affected and “unaffected” areas of the brain. These data have broad implications for interpreting transcriptomic data in neurodegenerative disorders.

---

Neurodegenerative proteinopathies such as Alzheimer’s disease (AD) and progressive supranuclear palsy (PSP) are characterized by aggregation and accumulation of self-proteins within insoluble aggregates^1^. AD is a complex proteinopathy characterized by extracellular amyloid β (Aβ) protein deposits and intracellular neurofibrillary tangles (NFTs) composed of the microtubule associated protein tau^2^. In PSP, which is considered a pure tauopathy, tau pathology is observed in several cell types. Tau accumulates as NFTs in neurons, as “tufts” in astrocytes (hence, the descriptor “tufted astrocytes”), and in coiled bodies or glial inclusions in oligodendrocytes^3^. In both diseases, numerous lines of research show a strong link between protein aggregation, accumulation and degeneration, though precise mechanism of cellular dysfunction and death remain enigmatic. Indeed, there is little consensus as to the mechanisms underlying cell dysfunction and death in AD, PSP and other neurodegenerative proteinopathies. Because of this incomplete understanding multiple studies are now using system level omics approaches to try and further understand the pathological cascades in AD, PSP and other neurodegenerative proteinopathies^4-6^. Here we compare the transcriptomic architecture in two brain regions from a large series of postmortem AD, PSP and control brains.

## Transcriptomic changes are conserved between AD and PSP

We compared the change in gene expression between AD and control and PSP and control in the temporal cortex (TCx) and cerebellar cortex (CER)^5,7^. Table 1A depicts the samples and data used. More extensive metadata, methodological details and raw RNAseq data can be found at AMP-AD Knowledge Portal (Supplementary Table S1). At a genome wide level, the data has been analyzed using two analytic models^5^. First, a simple model in which differential gene expression was conducted using linear regression with expression as the dependent variable, diagnosis as independent variable of primary interest, and RIN, age, sex, source of samples and flowcell as covariates (Supplementary Tables S2-S5). Second, a comprehensive model was applied to partially account for cell-type changes (Supplementary Tables S6-S9). The comprehensive model uses expression of five genes that serve as cell type markers (*ENO2* for neuron, *CD68* for microglia, *OLIG2* for oligodendrocyte, *GFAP* for astrocyte and *CD34* for endothelial cells) as covariates, in addition to all covariates in simple model^7^. For the analyses described here, we filtered the TCx and CER data for protein coding genes detected in both data sets above background based on their conditional quantile normalized values^5^. This filtering resulted in the identification of 14662 common genes in TCx and CER with associated β coefficients and q values of differential expression (DE) between AD and control and PSP and control. For comprehensive model analyses, this number is 14557 due to the exclusion of five cell type marker genes. Table 1B shows the summary data of the differentially expressed genes (DEGs), revealing large-scale transcriptomic changes in the protein-coding transcriptome for the AD TCx and CER with fewer DEGs withstanding false discovery in PSP. Using this data, we generated plots of the β coefficients of AD versus control (x-axis) and PSP versus control (y-axis) DE, using either no additional filter or filtering for various q value (i.e. false discovery rate adjusted p value) cutoffs. Even when examining all genes without a DEG q-value filter, there is a strong positive correlation between the changes observed in AD versus control and PSP versus control (Fig. 1A, B). Assessing data from the simple model for all genes using linear regression the R^2^ is 0.27 (Fig. 1A, p < 1.0e-10, slope 0.31) for TCx, and the R^2^ is 0.69 (Fig. 1B, p < 1.0e-10, slope 0.78) for CER. These R^2^ values are increased and remain highly significant when analyzed using the comprehensive model. In TCx the R^2^ is 0.62 (Fig. 1C, p < 1.0e-10, slope 0.85) and in CER the R^2^ is 0.39 (Fig 1D, p < 1.0e-10, slope 0.46). In either model, increasing the cutoff for q-value to 0.1, 0.05 or 0.01 reduces the number of genes but increases the strength of the correlations, with R^2^ ranging from 0.89 to 0.98 and slopes ranging from 0.77 to 1.13 (Fig. 1E).

**Table 1A.**
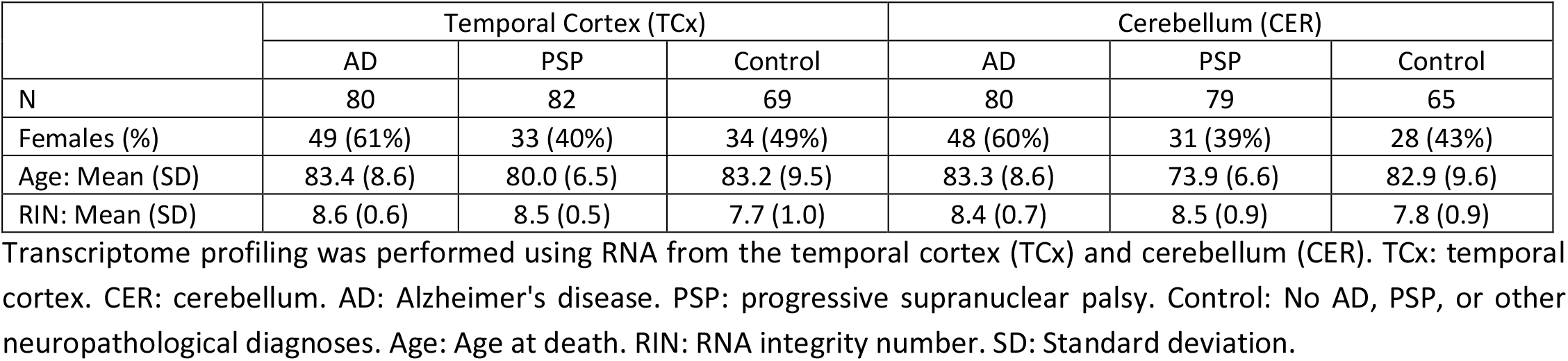
Characteristics of samples in the study

|  | Temporal Cortex (TCx) |  |  | Cerebellum (CER) |  |  |
| --- | --- | --- | --- | --- | --- | --- |
|  | AD | PSP | Control | AD | PSP | Control |
| N | 80 | 82 | 69 | 80 | 79 | 65 |
| Females (%) | 49 (61%) | 33 (40%) | 34 (49%) | 48 (60%) | 31 (39%) | 28 (43%) |
| Age: Mean (SD) | 83.4 (8.6) | 80.0 (6.5) | 83.2 (9.5) | 83.3 (8.6) | 73.9 (6.6) | 82.9 (9.6) |
| RIN: Mean (SD) | 8.6 (0.6) | 8.5 (0.5) | 7.7 (1.0) | 8.4 (0.7) | 8.5 (0.9) | 7.8 (0.9) |

Transcriptome profiling was performed using RNA from the temporal cortex (TCx) and cerebellum (CER). TCx: temporal cortex. CER: cerebellum. AD: Alzheimer's disease. PSP: progressive supranuclear palsy. Control: No AD, PSP, or other neuropathological diagnoses. Age: Age at death. RIN: RNA integrity number. SD: Standard deviation.

**Table 1B.**
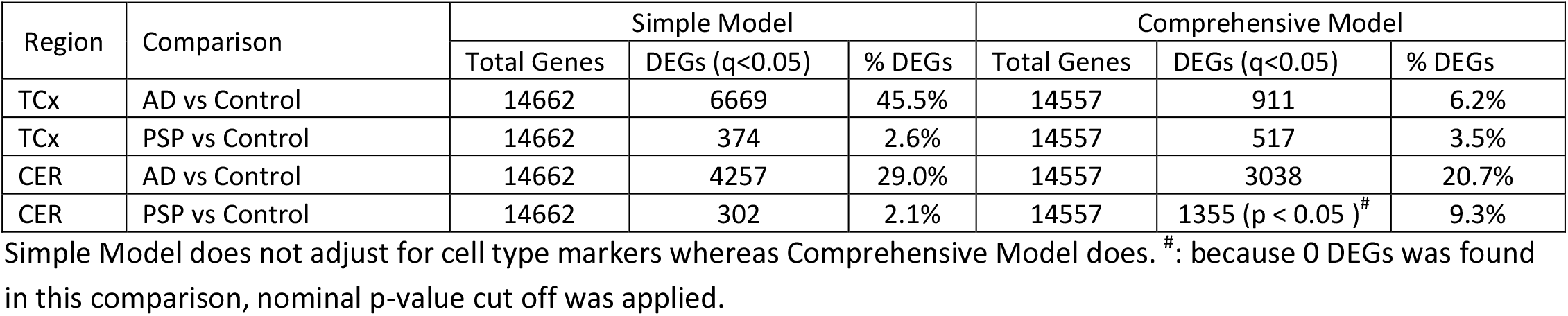
Summary of number of differentially expressed genes (DEGs)

| Region | Comparison | Simple Model |  |  | Comprehensive Model |  |  |
| --- | --- | --- | --- | --- | --- | --- | --- |
|  |  | Total Genes | DEGs (q<0.05) | % DEGs | Total Genes | DEGs (q<0.05) | % DEGs |
| TCx | AD vs Control | 14662 | 6669 | 45.5% | 14557 | 911 | 6.2% |
| TCx | PSP vs Control | 14662 | 374 | 2.6% | 14557 | 517 | 3.5% |
| CER | AD vs Control | 14662 | 4257 | 29.0% | 14557 | 3038 | 20.7% |
| CER | PSP vs Control | 14662 | 302 | 2.1% | 14557 | 1355 (p < 0.05 )<sup>#</sup> | 9.3% |

Simple Model does not adjust for cell type markers whereas Comprehensive Model does. <sup>#</sup>: because 0 DEGs was found in this comparison, nominal p-value cut off was applied.

**Figure 1.**
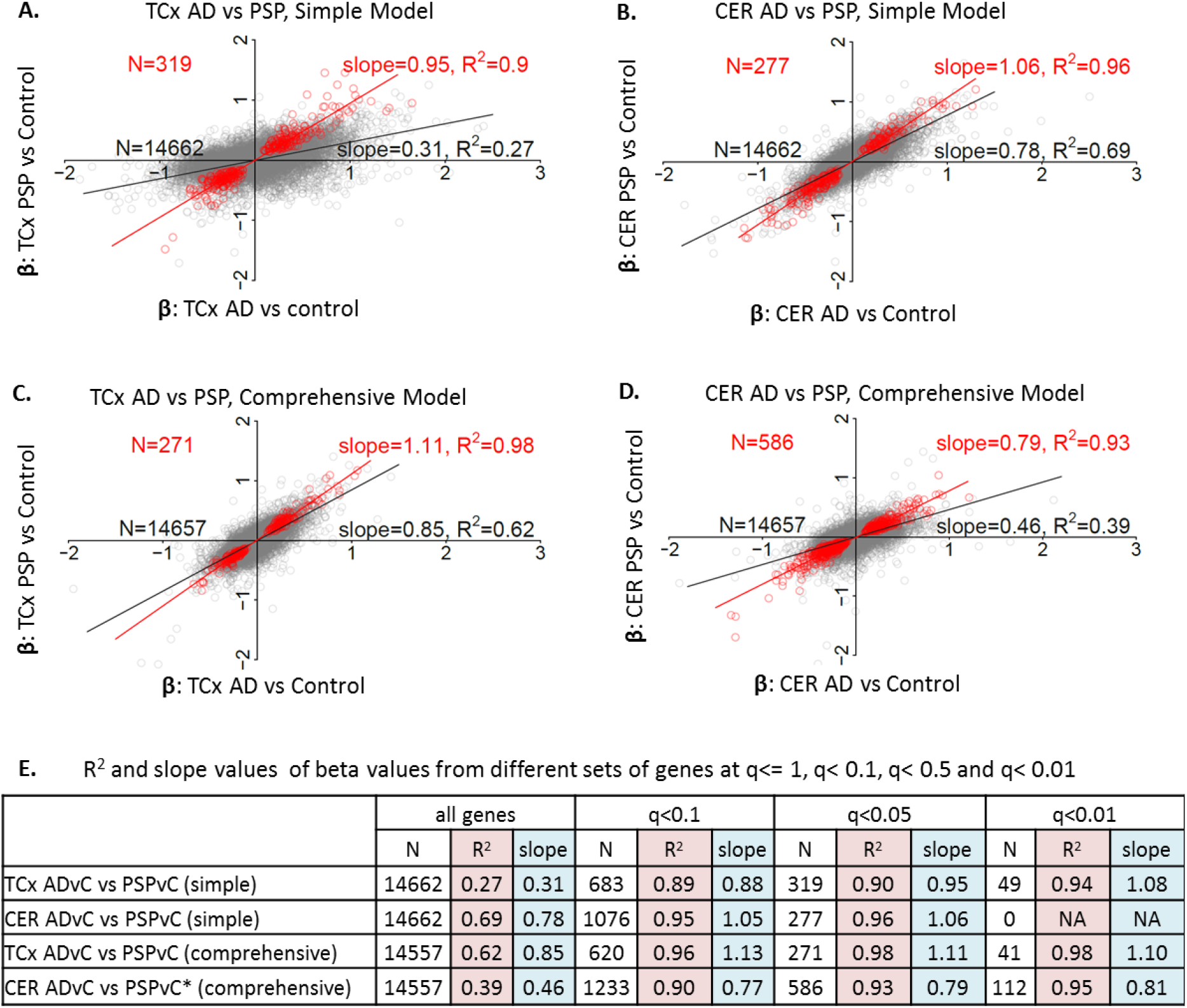
Gene expression changes are conserved between AD and PSP. **(A)-(D):** Comparison between beta coefficients (β) of AD vs control (ADvC) and those of PSP vs control (PSPvC) DEG analyses. Each circle represents a gene. Red circles: DEGs of q value < 0.05 on both side comparisons, except for (D) CER PSPvC where p value < 0.05 was used. Simple model: β is from linear regression with expression as dependent variable, diagnosis as independent variable of primary interest, and with RIN, age at death, sex, source of samples and flowcell as covariates. Comprehensive model: β is from linear regression as in simple model, with five additional covariates - expression of five cell type markers (*ENO2* for neuron, *CD68* for microglia, *OLIG2* for oligodendrocyte, *GFAP* for astrocyte and *CD34* for endothelial cells). **(E)** Summary of slope and R^2^ values between β of ADvC and those of PSPvC. *: p value cutoff instead of q value cutoff was applied when selecting DEGs in CER PSPvC comprehensive model.

These analyses show a striking conservation in the overall patterns of gene expression in two neurodegenerative disorders in two regions of the brain. These regions at the level of visible and gross pathologies are quite distinct. TCx is severely affected in AD^8^. It is atrophied with prominent neuronal synaptic loss and shows robust amyloid and tau pathologies and gliosis. In PSP, TCx tau pathology and neuronal loss is less severe than that in AD and even other regions of the brain affected earlier in the PSP disease course^9^. In contrast, CER is not typically reported to be pathologically affected in either AD or PSP, though certainly, in PSP deep cerebellar nuclei are affected. Nonetheless, connections between CER and brain areas may be damaged by both disorders^10^. Both the overall correlations in the entire set of genes analyzed and the increasing correlations observed when a q-value filter is applied, demonstrate that the transcriptomic architecture for protein coding genes is highly similar in these two disorders and that DEGs selected based on q-value cutoffs represent core transcriptome changes observed during neurodegeneration. Further, as bulk RNAseq data from whole brain tissue is strongly influenced by changes in cell type composition^11^, we note that the comprehensive model that takes into account these cell type changes shows a stronger correlation in the TCx between the disease states when compared to the simple model when no q-value cutoff is used. As the CER is relatively unaffected in terms of alterations in cell-type composition, when all genes are analyzed the correlation is actually weaker. Once a q-value filter is applied, there is little difference between the models. Such data indicates that cell-type changes indeed contribute to some of the transcriptome variance observed and correcting for that variance in the bulk RNAseq data can increase the power of the study to detect DEGs replicably across neurodegenerative diseases, when a tissue has cell type changes in one or both conditions, but may impair analyses when no large-scale cell type changes are presents.

## Transcriptomic changes are conserved across TCx and CER

The DEG changes between AD and PSP in two regions of the brain demonstrate a striking conservation of transcriptomic changes across these different neurodegenerative diseases. In designing these studies, we considered CER as an internal control for a relatively unaffected area of the brain. However, given the large number of highly significant DEGs in the AD CER, we evaluated whether the transcriptomic changes in the TCx and CER were also conserved within a disease classification (Fig. 2). In this case, we plotted the β coefficients for AD versus control in the TCx (x-axis) versus the β coefficients for AD versus control in the CER (y-axis) and likewise generated plots of the β coefficients of TCx versus CER for PSP versus control. Data from both the simple and comprehensive models are plotted. These analyses showed robust correlations. In AD, the overall R^2^ between TCx and CER was 0.35 (Fig. 2A p < 1.0e-10, slope 0.40) using the simple model and R^2^ = 0.32 (Fig. 2C, p < 1.0e-10, slope 0.63) using the comprehensive model. In PSP the overall R^2^ was 0.31 (Fig 2B, p < 1.0e-10, slope 0.59) in the simple model and R^2^ was 0.15 (Fig. 2D, p < 1.0e-10, slope 0.3) in the comprehensive model. Again, as the stringency of the q-value used to select the DEGs was increased both R^2^ (ranging from 0.70 to 0.95) and the slope (0.62 to 1.03) of the best-fit line increased when comparing the transcriptomes for the TCx and CER within disease states (Fig. 2E). Thus, not only is the transcriptomic architecture conserved between AD and PSP, it is also conserved across a severely affected and “unaffected” brain region in AD and a moderately affected and less affected brain region in PSP.

**Figure 2.**
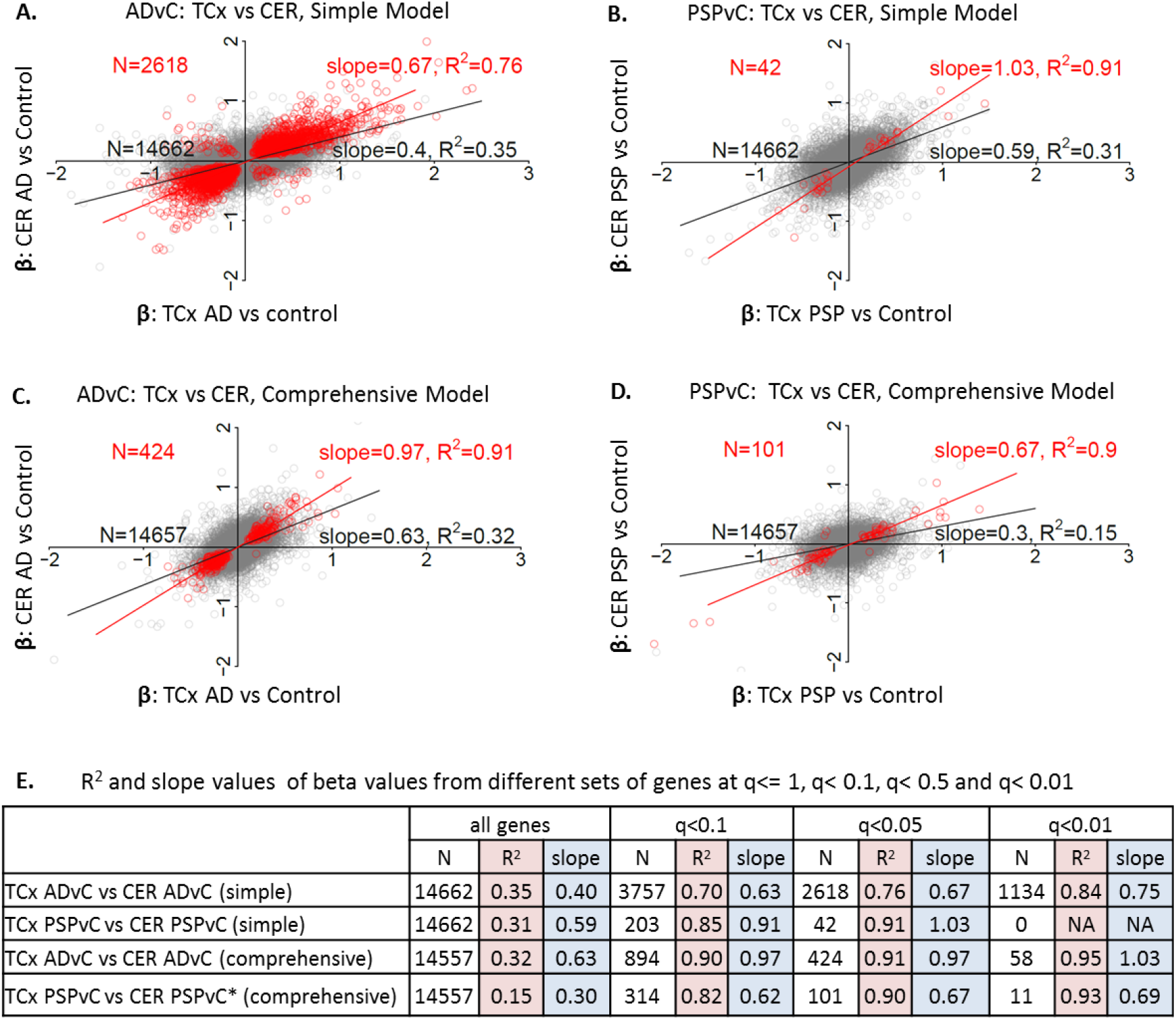
Gene expression changes are conserved between brain regions within disease analyses. **(A)-(D):** Comparison between beta coefficients (β) of TCx AD vs control (ADvC) and those of CER ADvC, and between TCX PSPvC and CER PSPvC DEG analyses. Each circle represents a gene. Red circles: DEGs of q value < 0.05 on both side comparisons, except for (D) PSPvC where p value < 0.05 was used. Simple model: β is from linear regression with expression as dependent variable, diagnosis as independent variable of primary interest, and with RIN, age at death, sex, source of samples and flowcell as covariates. Comprehensive model: β is from linear regression as in simple model, with five additional covariates - expression of five cell type markers (*ENO2* for neuron, *CD68* for microglia, *OLIG2* for oligodendrocyte, *GFAP* for astrocyte and *CD34* for endothelial cells). **(E)** Summary of slope and R^2^ values between β of TCx ADvC (or PSPvC) and those of CER ADvC (or PSPvC). *: p value cutoff instead of q value cutoff was applied when selecting DEGs in CER PSPvC comprehensive model.

## Gene Ontology Analyses

Given these striking correlations of DEG changes across two neurodegenerative disorders and two brain regions, we used gene ontology analyses to provide some biological context to these data. In this case, we binned the input into the GO analyses by focusing on DEGs (q value < 0.1) that were changed in the same direction. Thus, we first analyzed DEGs down in AD and PSP or up in AD and PSP using FUMA GWAS web server (https://fuma.ctglab.nl/)^12^. These data are summarized in Fig. 3 with more detailed versions provide in Supplementary Tables S10-S15. Shared upregulated DEGs in the TCx of AD and PSP are enriched (enrichment q value < 0.05) for biologic processes related to chromatin modification, gene expression, chromosome organization and metabolism of nucleotides. In the CER the shared upregulated genes link to biological processes relating to RNA and RNA transcription, cell-cell junctions, and heart, kidney, gland, and circulatory system development. Shared down regulated genes in AD and PSP are associated with GO cell compartment terms related to mitochondrial and ribosomal functions in both the TCx and the CER. These data and the extended GO analyses (Supplementary Tables S10-S15), point to highly complex biological changes shared in both AD and PSP.

**Figure 3.**
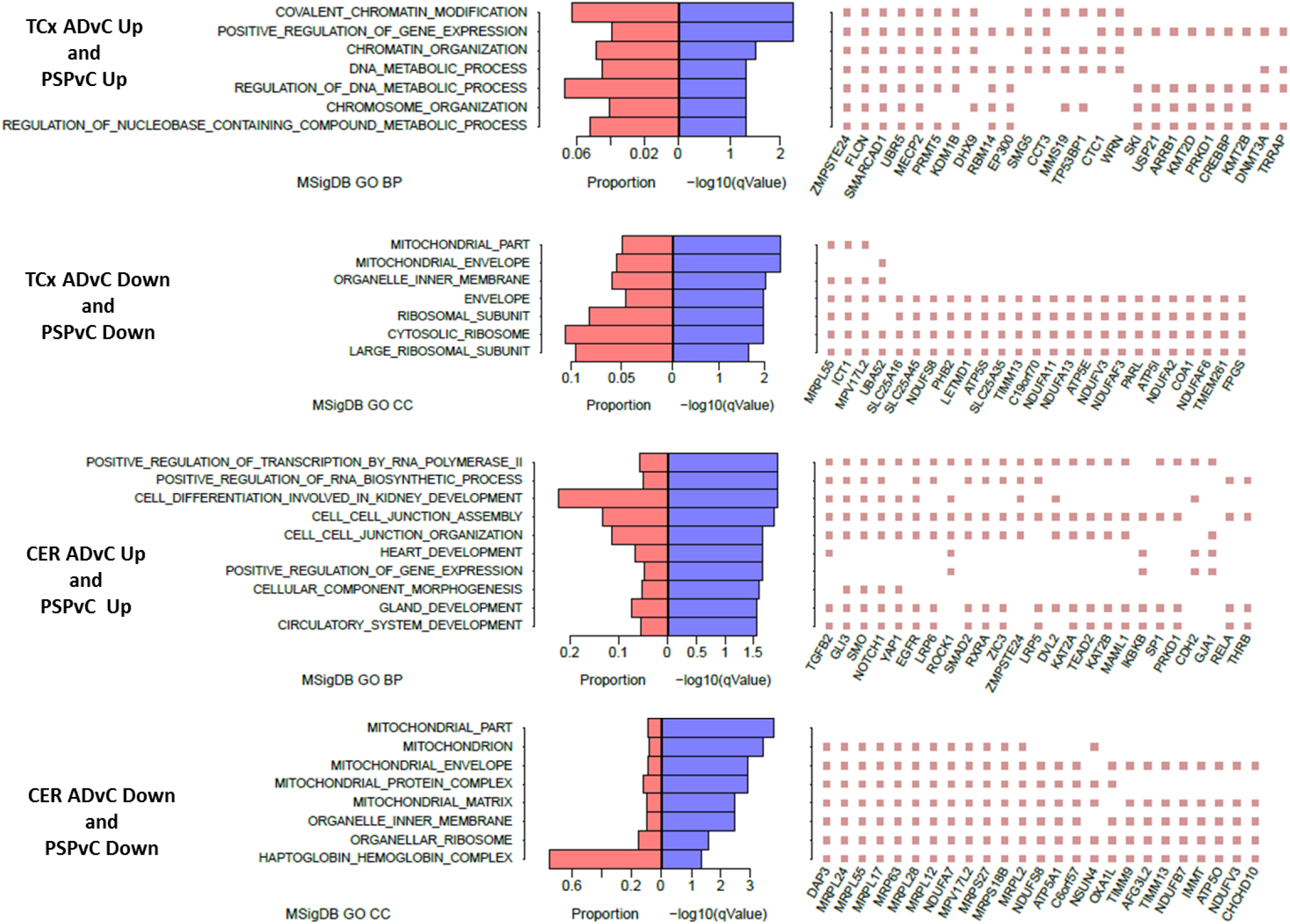
Gene ontology (GO) enrichment of differentially expressed genes (DEGs). Left panel: GO BP (biological process) terms of enrichment q-value < 0.05 were illustrated; when no such BP or molecular function term exists, CC (cellular compartment) terms of enrichment q-value < 0.05 were illustrated. Middle panel: -log10 enrichment q-value (blue bar) and proportion of DEGs in GO term over GO term genes (red bar). Right panel: top 25 DEGs that are mostly observed in selected GO terms. DEGs were identified at q<0.1 in both AD vs control and PSP vs control comparisons.

## Discussion

Numerous studies analyzing large-scale transcriptomic alterations in AD reveal a large number of network abnormalities that demonstrate widespread changes in pathways including but not limited to immune function, myelination, synaptic transmission and lipid metabolism^4,5,11,13-16^. Though these postmortem cross-sectional data sets provide a detailed systems level description of changes that have occurred over the disease course, in isolation they do not provide a framework for cause and effect relationships. The conservation in the overall transcriptome signature of AD and PSP relative to control brains indicates that the transcriptomic changes observed are more likely attributable to common downstream events in the neurodegenerative cascade and not initiating events. The fact that these conserved transcriptomic changes are observed in regions with neuropathologies varying from minimal to significant suggests that these conserved expression changes are unlikely to be driven by gross neuropathology or cell proportion changes. We have previously identified reduced expression of myelination network transcripts and proteins in both AD and PSP TCx and nominated it as a common disease mechanism for both conditions^5^. Given that AD and PSP are both tauopathies, conserved transcriptional alterations may not generalize to all neurodegenerative disorders. That said, the conservation holds in the CER, which is thought to be largely unaffected in these disorders, and therefore we would speculate that carefully conducted transcriptomic studies that are expanded to include other neurodegenerative proteinopathies may well show similar shared transcriptomic changes reflecting a long-standing neurodegenerative process triggered by protein accumulation.

Our finding that there is a shared transcriptomic architecture between the TCx and the CER within AD and PSP is noteworthy and consistent with our prior findings in transcriptional networks^5^. As noted previously, we had intended the CER to serve as a “control” for a largely pathologically unaffected brain region in AD; however, these transcriptomic data indicate a strong correlation between DEGs in both regions. Though this correlation is more robust due to the larger number of DEGs in AD vs. control, the correlation holds in PSP. This observation has several implications. First, these data demonstrate that long-standing neurodegenerative disease processes have a broad impact on the brain that extends well beyond visible pathology. Thus, there needs to be appropriate caution when inferring that a brain region in disease is “unaffected” based on an absence of pathological abnormalities as assessed using standard methods. Second, highly similar transcriptomic alterations in the brain driven by a regional or multi-regional proteinopathy likely reflect a mixture of common degenerative and compensatory responses attributable to long standing pathology within the brain, such as dysregulations of mitochondria^17^. Third, it is possible that the combination of epigenetic and genetic factors contributes to the similar transcriptomic alterations, as indicated by the DEGs of chromatin modification pathway^18,19^(Fig. 3).

In summary, the concept that AD, PSP or any other neurodegenerative disease has a specific transcriptomic signature may be inaccurate; rather there appears to be conserved transcriptomic alterations due to common proteinopathies or their downstream effects. This assertion will require additional large-scale transcriptomic analyses of other age associated neurodegenerative diseases conducted in a manner that eliminates many of the experimental confounds, such as batch effects. The large number of highly perturbed networks in AD that have been established in prior studies and our analyses in this study reinforce the notion that in the symptomatic phase, neurodegenerative diseases are characterized by incredibly complex biology that likely represents a mix of long-standing degenerative and compensatory processes. Such data reinforce the need to both develop paradigms that allow for the earliest possible intervention in these disorders that typically have long prodromal phases, and to develop multifaceted therapies that might be able to better alter the complex alterations present in the symptomatic phases of disease. Our findings also demonstrate the widespread perturbations of systems in the whole brain in neurodegenerative diseases, which requires novel biomarkers capable of tracking these changes in relatively “unaffected” brain regions and formulating therapies that address these ubiquitous alterations.

## Methods

### Subjects and Samples

The study dataset has been made available to the research community and described in detail previously^7,20^. Briefly, AD, PSP and control subjects were diagnosed neuropathologically at autopsy. AD subjects are from the Mayo Clinic Brain Bank, had definite neuropathologic diagnosis according to the NINCDS-ADRDA criteria^21^ and had Braak neurofibrillary tangle (NFT) stage of ≥4.0. All PSP subjects are from the Mayo Clinic Brain Bank and were diagnosed according to NINDS neuropathologic criteria^9^. Control subjects, either from Mayo Clinic Brain Bank or Banner Sun Health Institute, had Braak NFT stage of 3.0 or less, CERAD neuritic and cortical plaque densities of 0 (none) or 1 (sparse) and lacked various pathologic diagnoses. TCx and CER samples underwent RNA extractions via the Trizol/chloroform/ethanol method, followed by DNase and Cleanup of RNA using Qiagen RNeasy Mini Kit and Qiagen RNase -Free DNase Set. The quantity and quality of RNA samples were determined by the Agilent 2100 Bioanalyzer using the Agilent RNA 6000 Nano Chip. Samples included in this study all have RIN ≥5.0. Among final samples included in this study (231 TCx samples and 224 CER samples), 197 TCx and 197 CER samples were paired, i.e. from the same 197 subjects.

### RNA sequencing

Library preparation and sequencing of the samples were conducted at the Mayo Clinic Genome Analysis Core using TruSeq RNA Sample Prep Kit (Illumina, San Diego, CA). The library was sequenced on Illumina HiSeq2000 instruments, generating 101 base-pair, paired-end raw reads. Raw reads were processed through MAPR-Seq pipeline^22^ v1.0 which removed reads of low base-calling Phred scores, aligned remaining reads to human reference genome build GRCh37 using Tophat v2.0.12^23,24^, counted reads in genes using Subread 1.4.4^25^, and obtained QC measures from both pre-alignment reads and post-alignment reads using RSeQC toolkit^26,27^ and fastQC (https://www.bioinformatics.babraham.ac.uk/projects/fastqc/). Samples that have high RNA degradation, or low reads mappability, or inconsistency between recorded sex and estimated sex using RNAseq chromosome Y expression were removed from downstream analysis. Raw reads of remaining samples were normalized using R cqn package^28^ which took into consideration library size, gene GC content and gene coding length, resulting in normalized expression in log2 scale. Additional information could be found in our previous publication^5,20^.

### Regression analysis

Multiple linear regression (MLR) were performed for each gene using normalized gene expression as dependent variable, diagnosis as primary independent variable, and RIN, age at death, sex, source of samples and flowcell as covariates (simple model), plus expression of five cell type markers (*ENO2* for neuron, *CD68* for microglia, *OLIG2* for oligodendrocyte, *GFAP* for astrocyte and *CD34* for endothelial cells) as covariates (comprehensive model), as previously published^5^. Diagnosis groups in these MLR were TCx ADvC, TCx PSPvC, CER ADvC and CER PSPvC.

Using β coefficients of DEGs of q-value<=1 (namely, <0.1, 0.05, 0.01) from the above MLR, simple linear regression was performed. Slopes and R^2^ were obtained (Fig 1-2) from the following models: β.TCx.PSPvsCtrl ∼ 1 + slope * β.TCx.ADvsCtrl, β.CER.PSPvsCtrl ∼ 1 + slope * β.CER.ADvsCtrl, β.CER.ADvsCtrl ∼ 1 + slope * β.TCx.ADvsCtrl, and β.CER.PSPvsCtrl ∼ 1 + slope * β.TCx.PSPvsCtrl.

### GO enrichment analysis

Differentially expressed genes were analyzed for GO enrichment using FUMA GWAS web server at https://fuma.ctglab.nl with MSigDB v7.0 ^12,29^. Background genes (N=14662) were the expressed coding genes in both TCx and CER cohorts, genes of interest were DEGs of q-value < 0.1 in both group comparisons and consistent in direction of expression change. Figures were made using R software environment.

## Data Availability

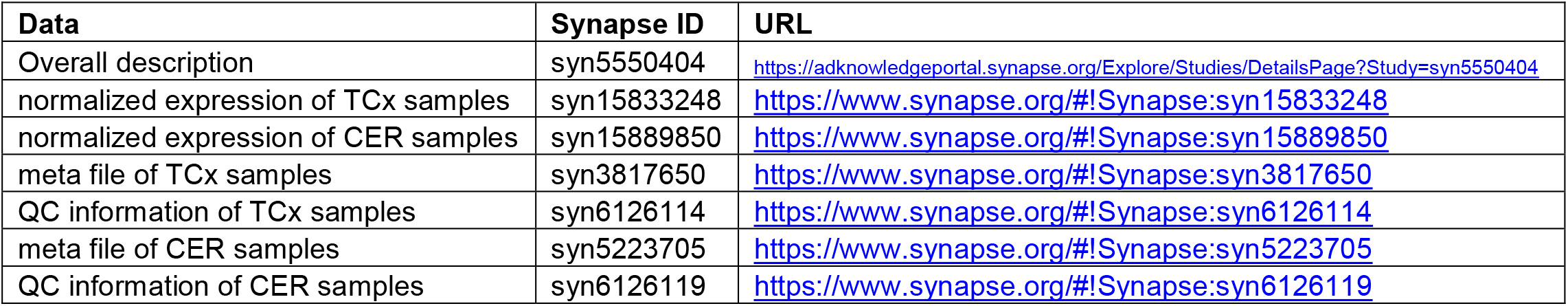

Data Availability:
| Data | Synapse ID | URL |
| --- | --- | --- |
| Overall description | syn5550404 | https://adknowledgeportal.synapse.org/Explore/Studies/DetailsPage?Study=syn5550404 |
| normalized expression of TCx samples | syn15833248 | https://www.synapse.org/#!/Synapse:syn15833248 |
| normalized expression of CER samples | syn15889850 | https://www.synapse.org/#!/Synapse:syn15889850 |
| meta file of TCx samples | syn3817650 | https://www.synapse.org/#!/Synapse:syn3817650 |
| QC information of TCx samples | syn6126114 | https://www.synapse.org/#!/Synapse:syn6126114 |
| meta file of CER samples | syn5223705 | https://www.synapse.org/#!/Synapse:syn5223705 |
| QC information of CER samples | syn6126119 | https://www.synapse.org/#!/Synapse:syn6126119 |

## Acknowledgments

We thank the patients and their families for their participation, without whom these studies would not have been possible.

Supported by the NIHNIA AMPAD U01 AG046139 (NET, NP, TEG,CF) and P30AG066506/P50AG047266 (TEG). The results published here are in whole or in part based on data obtained from the AD Knowledge Portal (https://adknowledgeportal.synapse.org/). Study data were provided by the following sources: The Mayo Clinic Alzheimers Disease Genetic Studies, led by Dr. Nilufer Ertekin-Taner and Dr. Steven G. Younkin, Mayo Clinic, Jacksonville, FL using samples from the Mayo Clinic Study of Aging, the Mayo Clinic Alzheimer’s Disease Research Center, and the Mayo Clinic Brain Bank. Data collection was supported through funding by NIA grants P50 AG016574, R01 AG032990, U01 AG046139, R01 AG018023, U01 AG006576, U01 AG006786, R01 AG025711, R01 AG017216, R01 AG003949, NINDS grant R01 NS080820, CurePSP Foundation, and support from Mayo Foundation. Study data includes samples collected through the Sun Health Research Institute Brain and Body Donation Program of Sun City, Arizona. The Brain and Body Donation Program is supported by the National Institute of Neurological Disorders and Stroke (U24 NS072026 National Brain and Tissue Resource for Parkinson’s Disease and Related Disorders), the National Institute on Aging (P30 AG19610 Arizona Alzheimer’s Disease Core Center), the Arizona Department of Health Services (contract 211002, Arizona Alzheimer’s Research Center), the Arizona Biomedical Research Commission (contracts 4001, 0011, 05-901 and 1001 to the Arizona Parkinson’s Disease Consortium) and the Michael J. Fox Foundation for Parkinson Research.

